# Electronic Polarizability Tunes the Function of the Human Bestrophin 1 Cl⁻ Channel

**DOI:** 10.1101/2023.11.14.567055

**Authors:** Linda X. Phan, Aaron P. Owji, Tingting Yang, Jason Crain, Mark S.P. Sansom, Stephen J. Tucker

## Abstract

Mechanisms of anion permeation within ion channels and nanopores remain poorly understood. Recent cryo-electron microscopy structures of the human bestrophin 1 Cl⁻ channel (hBest1) provide an opportunity to evaluate ion interactions predicted by molecular dynamics (MD) simulations against experimental observations. Here, we implement the fully polarizable forcefield AMOEBA in MD simulations on different conformations of hBest1. This forcefield models multipole moments up to the quadrupole; therefore, it captures induced dipole and anion-π interactions. We show that key biophysical properties of the channel can only be simulated when electronic polarization is included in the molecular models and that Cl⁻ permeation through the neck of the pore is achieved through hydrophobic solvation concomitant with partial ion dehydration. Furthermore, we demonstrate how such polarizable simulations can help determine the identity of ion-like densities within high-resolution cryo-EM structures and that neglecting polarization places Cl⁻ at positions that do not correspond with their experimentally resolved location. Overall, our results demonstrate the importance of including electronic polarization in realistic and physically accurate models of biological systems, especially channels and pores that selectively permeate anions.

**Statement of Significance:** Ion channels are nanoscale protein pores that enable the selective passage of charged ions across cell membranes. Understanding the underlying mechanisms for selective anion permeation through such pores remains a challenge. To simulate their behavior efficiently *in silico*, fixed charge models are typically employed. However, this approach is insufficient for the study of anions. Here, we use simulations with explicit treatment of electrostatics to investigate the interactions of chloride ions in the human bestrophin 1 channel. We find that electronic polarization tunes the state of the channel and affects the interactions of chloride ions thereby revealing a mechanism for permeation. Furthermore, these simulations can be used to resolve experimental ambiguity in ion-like densities from cryo-EM structures.

## Introduction

Ion channels are nanoscale pore-forming membrane proteins that enable the rapid and selective passage of ions across a membrane. Their activity and function are central to the regulation of physiological processes from cell signaling and the control of pH balance, to muscle contraction and brain function ^1,2^. Chloride ions are the most abundant anion in living organisms. Since dysfunction of associated Cl⁻ channels is known to result in a variety of disease states, these proteins represent attractive therapeutic targets ^3^. However, the underlying mechanisms of Cl⁻ permeation and selectivity in these channels remain under-explored relative to their cation counterparts, and is partly a consequence of their weak selectivity ^4^. It is therefore of great interest to explore the mechanisms of anion permeation in such channels and to thoroughly assess the essential physics needed for accurate functional annotation.

Studies suggest that Cl⁻ can form favorable interactions with hydrophobic interfaces ranging from simple air/water interfaces to more complex protein interfaces ^5–7^. The anisotropy of such interfaces induces a dipole in the Cl⁻ that is otherwise not present in bulk. Interactions between the induced dipole and surrounding water molecules compensate for the partial dehydration of Cl⁻ as it adsorbs at the interfacial layer and comes into direct contact with the hydrophobic interface ^8^. This phenomenon can be observed across the Hofmeister series i.e., F^-^ < Cl^-^ < Br^-^ < I^-^, whereby the softer, more polarizable anions are more prone to dehydration and localize at the interface ^9^.

Anion interactions with aromatic edges are also a well-established phenomenon ^10–12^ and more recently been recognized to play important roles in buried regions of proteins ^11,13^. Also, the ion conduction pathway within some anion channels can be composed of aromatic residues such as phenylalanine. Known structures of anion channels that exploit this include the mechanosensitive Cl^-^ channel, Flycatcher1 (FLYC1) where the narrowest constriction (∼ 2.8 Å radius) is created by a ring of phenylalanine side chains ^14^, the slow anion channel (SLAC1) which is occluded by a highly conserved phenylalanine residue responsible for channel gating (< 2 Å) ^15^ and the mechanosensitive channel of small conductance from *Escherichia coli* (EcMscS) in which phenylalanines appear to constitute the hydrophobic gate (∼ 4 Å) ^16^. However, although such anion-aromatic interactions are readily observed, the degree to which dilation is required for anion passage in the presence of pore-ling phenylalanine residues is variable and only partially understood. Therefore, there is a need for further biophysical characterization to better understand the functional roles of such pore-lining aromatics and their contributions towards anion permeation.

Cryo-electron microscopy (cryo-EM) is a powerful structural technique that has revolutionized the field of structural biology. Recent technological improvements in sample treatment, grid preparation, microscope hardware, and image processing have made it achievable to obtain high-resolution maps better than 3 Å ^17^. Despite these advancements, there are several caveats, such as radiation damage, resulting in lower signal-to-noise ratios, protein denaturation and beam-induced sample movement, which all present limitations to achieving even higher resolutions ^18^. The resolution of single-particle cryo-EM is also often insufficient to provide detailed information on small molecules or bound ligands, such as water or ions, which require a resolution of at least 2.5 Å ^19^, and so it can be difficult to differentiate and interpret these small densities with confidence when using this technique. Molecular dynamics (MD) simulations can extend the capabilities of cryo-EM by capturing conformational variability and short-lived states. Furthermore, MD simulations can assist in the assignment and interpretation of cryo-EM data to access improved atomic resolution structures ^20^.

The majority of MD simulations employ pairwise additive forcefields that model electrostatic interactions as Coulombic forces between fixed point charges ^21^. These forcefields do not capture the effects of induced polarization that arise from redistributions of charge density in response to local field gradients. This becomes problematic for modelling systems that involve polarizable moieties, such as aromatic residues and polarizable anions. Charge-scaling methods, such as the prosECCo75 ^22–24^ and NBFIX ^25,26^ approaches, applied to existing pairwise additive forcefields, aim to improve molecular interactions by more accurately capturing the effects of induced polarization, leading to better agreement with experimental data. Furthermore, a number of biomolecular forcefields have recently been developed to explicitly capture induced polarization. Among these, two of the most widely employed are the AMOEBA forcefield ^27^ which models atomic monopoles through to quadrupole moments within a classical MD framework, and the CHARMM Drude forcefield ^28,29^, which uses massless Drude oscillators to displace charge from atomic centers. These forcefields provide new and improved levels of predictive power over non-polarizable forcefields and are increasingly being used to investigate proteins ^6,30,31^; however, they incur additional computational costs.

An example where modelling polarization may be vital is in the study of Cl⁻-selective ion channels. Due to the weakly polarizable nature of Cl⁻ ^32^, realistic behavior is subtle and often difficult to capture. A suitable model are the Bestrophin channels, a family of calcium-activated Cl⁻ channels (CaCCs) where many questions still remain ^33,34^. Four paralogs (Best1-4) have been identified in eukaryotes which are responsible for a diverse range of functions^34^. The best-known physiological role of Best1 is in the eye where disease-causing mutations cause retinal degenerative disorders called bestrophinopathies. The structures of bestrophin channels contain two key constrictions in the ion conduction pathway. A permeating ion from the extracellular side will first encounter the neck region, which is a gate composed of three hydrophobic residues (I76, F80 and F84) that are highly conserved across homologs. After passing this gate, the ion will traverse the interior of the cytosolic vestibule, followed by a second, shorter constriction at the cytosolic exit, called the aperture, which displays significant divergence across paralogs ^34^, but together, the neck and aperture are both thought to comprise part of the gating mechanism for bestrophin channels, although their relative contribution to gating and/or Cl⁻ selectivity remain unclear ^34–37^.

Comparison of the human Bestrophin 1 channel (hBest1) in a fully open state (PDB ID 8D1O, 2.4 Å resolution) (**Figure 1A-C, G**) with a “partially open neck” conformation (PDB ID 8D1K, 2.3 Å resolution) (**Figure 1D-G**) ^36^ reveals a conformational change in the pore-facing residues of the hydrophobic neck (**Figure 1B & 1E).** In particular, the three states differ in the radius profile in the neck of the pore, with minimum HOLE radii of 0.5 Å in the closed state, 4.5 Å in the open state and 2 Å in the 8D1K structure ^36^. The 8D1K structure has therefore been (provisionally) designated as “partially open” given that the radius of a fully hydrated chloride ion ∼ 4 Å ^38^ and we refer to this as the partially open state in this paper. This difference allows us to probe the mechanisms of ion permeation and selectivity, as well as the conformational pathway to channel opening.

**Figure 1:**
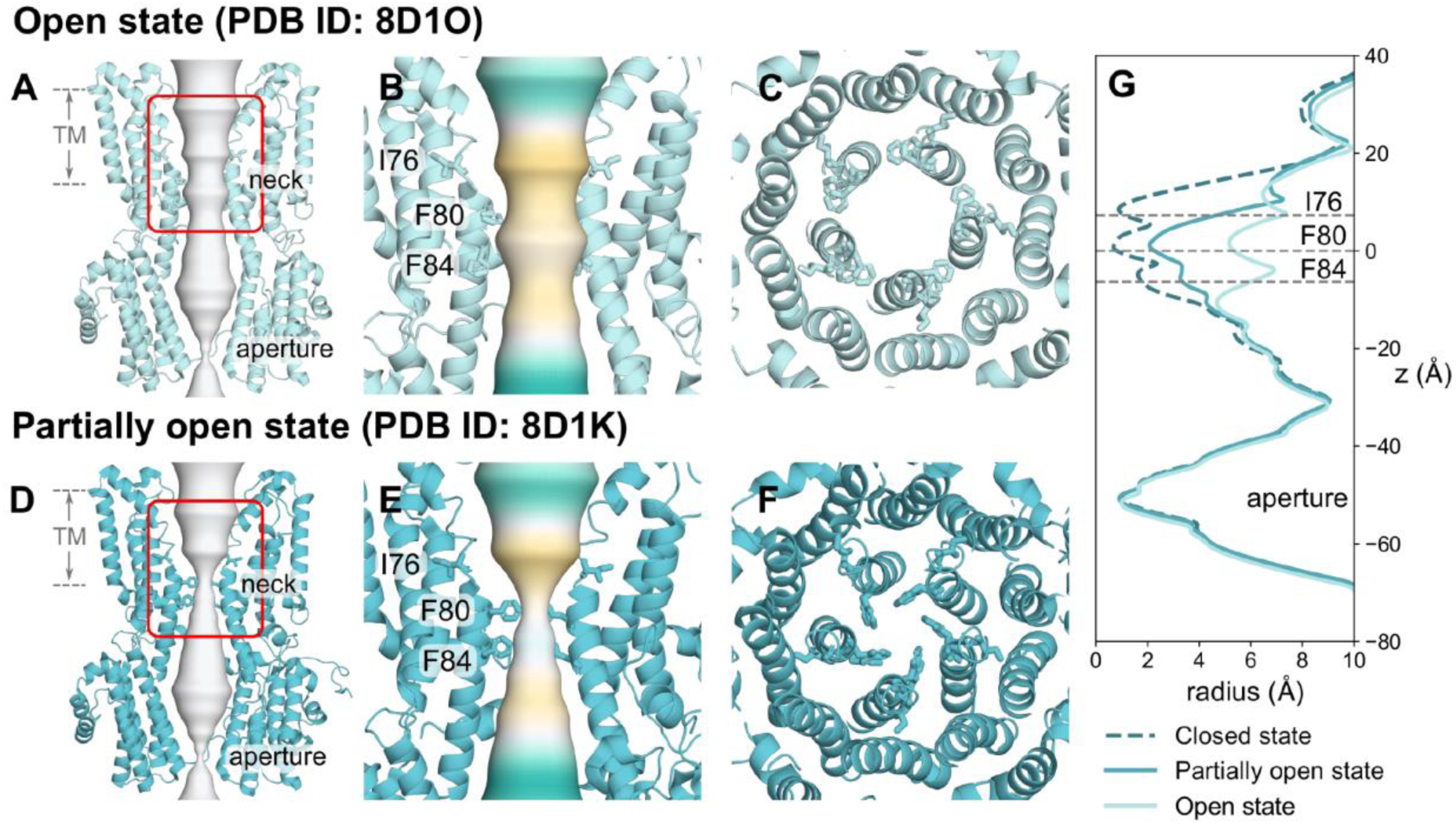
Cryo-EM structures of hBest1 in the open and “partially open” states. **A** Open state hBest1 (PDB ID 8D1O) and **D** partially open state hBest1 (PDB ID 8D1K) structures visualized with ion permeation pathway in white. The transmembrane neck region of the **B** open state and **E** partially open state structures, with ion permeation pathway colored by hydrophobicity with pale brown corresponding to maximum hydrophobicity and green corresponding to maximum hydrophilicity. Top-down view of the **C** open state neck and **F** partially open state neck. **G** Pore radius profiles of the open, partially open and closed (PDB ID 8D1I) states.

In this study, we therefore explore the effect of electronic polarization on hBest1 function by implementing fully polarizable forcefields to examine the behavior of Cl⁻ within the neck region. Here, we present an example of how such forcefields provide novel detailed insights into channel function and the behavior of Cl⁻ in the neck of hBest1. They also reveal the roles of aromaticity and pore asymmetry in ion permeation. In particular, phenylalanine sidechains are seen to undergo a conformational change which enables edge-on interactions with a partially dehydrated Cl⁻ during permeation. Crucially, we also show that models which include realistic polarization can be used to interpret ambiguous ion-like densities observed in experimental high-resolution cryo-EM structures, thus enabling more accurate functional annotation.

## Methods

### Structural model and system preparation

Cryo-EM structures of human Bestrophin 1 (hBest1) in the Ca^2+^–bound open state (PDB ID 8D1O, 2.4 Å resolution) and the hBest1 Ca^2+^–bound partially open neck state (PDB ID 8D1K, 2.3 Å resolution) ^36^ obtained from the Protein Data Bank (PDB). Due to the methodological and high computational demands for implementing explicitly polarizable forcefields, we reduced the protein structures to include the pore-lining segments spanning most of the transmembrane domain. The protein fragments were composed of residues 56 to 99, which encompass the neck region of interest (residues I76, F80 and F84) (**Figure S1**). Protein fragment systems were solvated in a 0.5 M NaCl solution and were prepared using the CHARMM-GUI protocol ^39^.

### Non-polarizable molecular dynamics

MD simulations employing a non-polarizable forcefield were performed in GROMACS ^40^ version 2021 using the CHARMM36m (c36m) forcefield in conjunction with the mTIP3P water model. Protein fragment systems were first subjected to energy minimization followed by a 2 ns NVT equilibration period. Simulations in the NPT ensemble were conducted for 100 ns whereby the first 20 ns of each simulation were discarded as an equilibration. Therefore, the final 80 ns of the simulation were used for analysis.

Simulations were carried out using the leap-frog integrator with a timestep of 2 fs. The temperature was maintained at 310 K using the Nosé-Hoover thermostat ^41^ with a time coupling constant of 1.0 ps. The pressure was maintained at 1 bar using the Parrinello-Rahman barostat ^42^ with a time coupling of 5.0 ps. Short-range electrostatics were treated with the Verlet cutoff scheme with a cutoff at 1.2 nm and long-range electrostatics were treated with the particle mesh Ewald (PME) algorithm ^43^. The LINCS algorithm ^44^ was used to constrain h-bonds. Backbone atoms were placed under harmonic restraints with a force constant of 1000 kJ/mol/nm^2^ to prevent the structures from deviating too much from the experimental coordinates. Three independent repeats were carried out for each system.

### Polarizable simulations molecular dynamics

Fully polarizable forcefield simulations were carried out in OpenMM 7.4.2 (www.openmm.org) and all components in the system were modelled with the AMOEBA polarizable forcefield using the amoeba2013 parameter set ^45^. Starting configurations for these simulations were obtained from the end of the NVT ensemble equilibration period using c36m as described above. Simulations were then setup following a similar procedure to a method described previously (https://github.com/Inniag/openmm-scripts-amoeba) ^6^.

AMOEBA forcefield simulations proceeded by performing 1000 steps of energy minimization to resolve any divergent energies due to the induced dipoles. The production run was simulated for 60 ns, with the first 10 ns of the simulation discarded for equilibration; therefore, the final 50 ns were used for analysis. Time integration was performed using the r-RESPA multiple time step integration algorithm ^46^ with an inner time step of 0.25 fs and an outer time step of 2 fs. The temperature was maintained at 310 K using the Andersen thermostat and pressure was maintained at 1 bar using the isotropic Monte Carlo barostat. Electrostatic multipole interactions were evaluated by the PME method with a real-space cutoff of 8 Å and tolerance of 5 × 10^−4^ and a fifth order B-spline interpolation. VdW interactions were calculated explicitly up to a distance of 12 Å and interactions beyond this cutoff treated with an analytical long-range dispersion correction. All C_α_ atoms were placed under a harmonic restraint with force constant 1000 kJ/mol/nm^2^ to prevent the protein from deviating from the experimental structure.

### Binding site identification and analysis

Interactions of Cl⁻ with each protein structure, binding site detection and quantification were calculated using PylipID (https://github.com/wlsong/PyLipID) ^47^. Binding sites were defined by Cl⁻ contact with three or more residues and by a dual-cutoff scheme for which Cl⁻ were considered bound if they resided between 3.3 Å and 4.8 Å based on data from the first hydration shell of Cl⁻ using the AMOEBA forcefield. Alignment and visualization of structures were achieved using Pymol (https://pymol.org/2/). Trajectory analysis was performed using MDAnalysis ^48,49^ and GROMACS analysis tools ^50^. The pore radius profiles were obtained using CHAP (www.channotation.org) ^51^.

### Results & Discussion

To probe the role of the conserved neck region in bestrophin channels, we have performed atomistic MD simulations of fragments from hBest1 containing the neck region and pore-lining sections from both the fully open (PDB ID 8D1O) and partially open neck states (PDB ID 8D1K). These fragments consisted of residues 56 to 99 from the full protein and were simulated in a 0.5 M NaCl solution (**Figure S1**). Previous studies of reduced systems of ion channels focusing on isolated sections have shown that they can provide an accurate representation of the dynamics within the original protein ^6,52,53^. To validate the protein fragment, we performed simulations of the full protein embedded in a lipid bilayer compared with the protein fragment and analyzed the average water density. The resulting profiles are similar, and the average water density remains close to bulk outside the neck region (**Figure S2**). This provides confidence that the protein fragments in our reduced system offer a representative model of the fully open and hydrated pore from the full protein.

The system contains aromatic residues and polarizable ions and therefore justifies the use of fully polarizable forcefields, such as AMOEBA. However, with added complexity comes increased computational costs, hence the need to reduce the system size to make the simulations computationally feasible.

### Interpretation of cryo-EM densities from fully polarizable MD simulations

In the experimental structure of the open state hBest1, a number of non-protein, ion-like densities have been observed, which have been modelled as water molecules in the published structure. A set of these water molecules (corresponding to HOH739 and HOH539 in the PDB) are located within the neck region and appear to bind to the helix dipole ^54^ at a distance of 3.2 Å to the backbone NH of F80 of each chain (**Figure 2A**). Similar non-protein densities are also observed consistently in proximity to the helix dipole in the open state structure of human bestrophin 2 (hBest2) (PDB ID 8D1N). Due to experimental limitations, there is ambiguity in the molecular identity of these densities. Assessment of the local chemistry and comparisons with known bromide binding sites from anomalous scattering studies of a Best1 homolog ^55^ suggest the location of these densities are comparable to the putative water molecules in the hBest1 structure. Therefore, it has been suggested that this site could possibly serve as a Cl⁻ binding site. Here, we have used the AMOEBA forcefield to explore the possibility that these densities could instead be representative of Cl⁻. We refer to the density X739 as the ion-like density previously labelled as a water molecule (HOH739) in the PDB.

**Figure 2:**
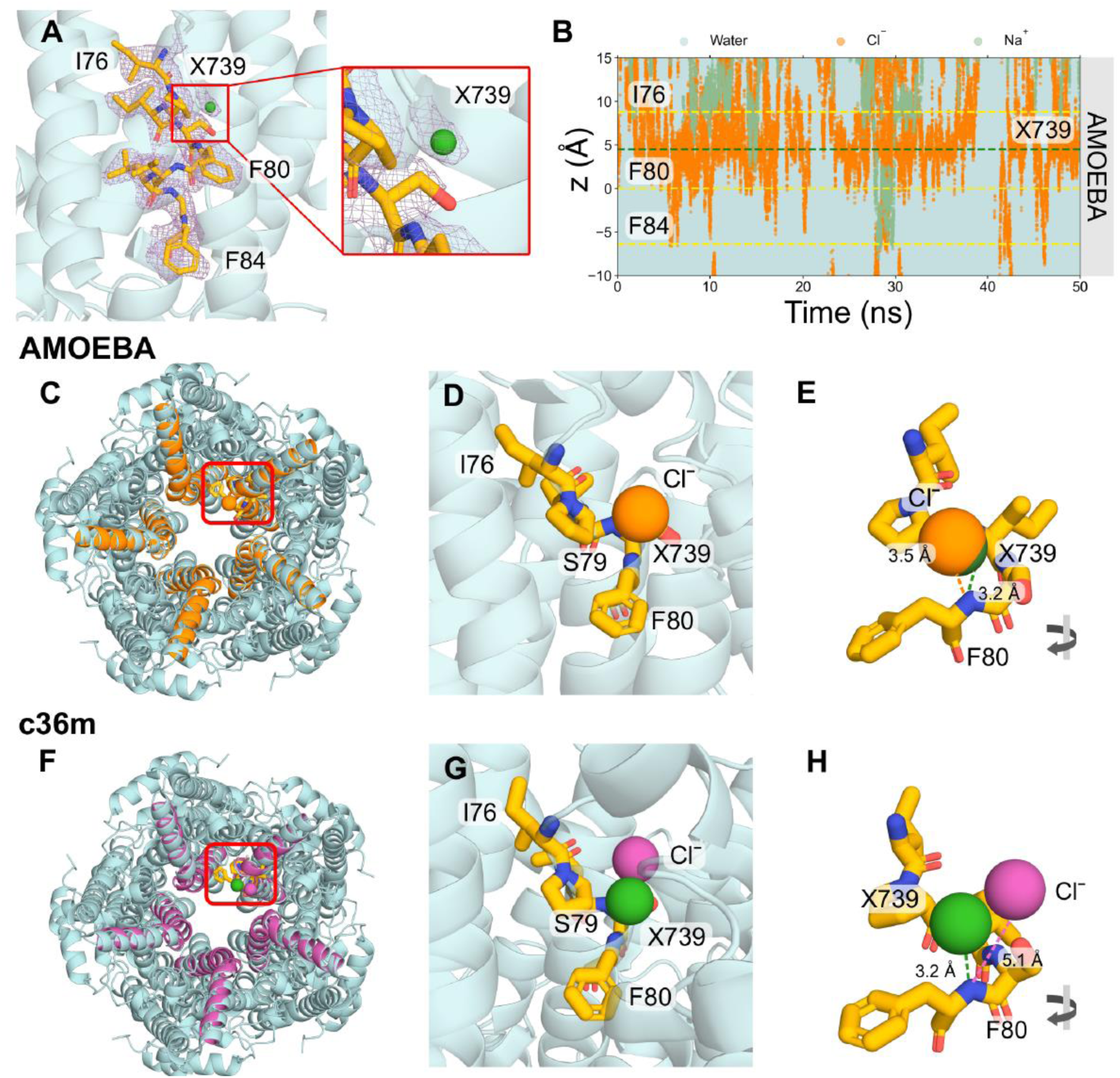
Cl⁻ binding site analysis. **A** Close-up representation of neck residues (I76, F80 and F84) from a single chain highlighted in licorice representation of the open state structure (PDB ID 8D1O). The cryo-EM densities are shown as a mesh representation in purple. A panel is zoomed in on the non-protein ion-like density in proximity to F80 labelled as X739, which corresponds to the water molecule HOH739 in the PDB. **B** The trajectories of water (cyan), Na⁺ (green) and Cl⁻ (orange) ions in z-coordinates as a function of time within the neck region of the pore. The plot indicates that Cl⁻ tends to cluster in a region between I76 and F80. The green dashed line represents the location of X739 from the experimental structure. AMOEBA forcefield simulations: Top-down view of the protein fragment in **C** AMOEBA (orange) or **F** c36m (magenta) aligned to the full protein structure (cyan). The location of X739 is indicated by the green sphere. The top detected binding pose identified with PyLipID shows **D** significant overlap between the AMOEBA Cl⁻ (orange sphere) and X739, indicating this site likely functions as a Cl⁻ binding site or **G** the c36m Cl⁻ (magenta sphere) shares no overlap with X739. **E, H** A rotated view showing the distance of the Cl⁻ and X739 to the backbone N of F80 in the experimental structure.

Analysis of the z-positions of ions and water as a function of time revealed that the open state neck is wetted and permeable to ions. There is a clear accumulation of Cl⁻ in the region between I76 and F80 of the neck (indicated by the dense orange region in **Figure 2B**). Na⁺ can be observed to be largely excluded from the neck region however a small fraction may traverse the neck (e.g. **Figure 2B** at ∼ 28 ns). Our simulations do not provide sufficient sampling for studying ion conductance ratios. To pinpoint precisely where Cl⁻ is clustering in three dimensions, we have used the computational tool, PyLipID, which is capable of analyzing protein-ligand interactions ^47^. Taking the top-ranked binding pose for the detected binding site, we focus on interactions of a single chain and have aligned the simulated fragment structures with the original PDB structure (chain E and HOH739) for comparison (**Figure 2C**).

The results of the AMOEBA simulations correlate very well with the cryo-EM densities. A number of identified binding sites in the neck region are within 0.5 Å of the ion-like density (**Figure S3**) and we show the binding site that is most representative of the density associated with HOH739 in the original structure (**Figure 2C-E**). A significant overlap can be seen between the Cl⁻ and the water molecule modelled in the PDB structure (**Figure 2D**). The distance of the Cl⁻ to the backbone nitrogen of F80 is 3.5 Å compared with 3.2 Å in the experimental structure (**Figure 2E**) and it is equidistant from the pore axis. Studies of CLC channels have revealed that pore-lining backbone amides can influence ion selectivity and permeation ^4^. Therefore, it is feasible that the ion-like density in this experimental structure is not water but a Cl⁻. This difference in ion location between the simulation and structural densities could be a result of the imposition of 5-fold symmetry during structural processing, which could marginally shift the cryo-EM density in comparison to allowed asymmetric interaction modes. Conversely, from a simulation perspective, the localization of Cl⁻ can be forcefield dependent and conditional on the realism of the model.

To demonstrate the impact of modelling polarizability, equivalent simulations were also performed with the non-polarizable forcefield, CHARMM36m (c36m) (**Figure 2F**). In contrast to the AMOEBA simulations, these simulations suggest that Cl⁻ tends to localize at a distance of 5.1 Å from the backbone nitrogen of F80 (**Figure 2H**) and prefers to interact with the backbone NH, hydroxyl group of S79, and the backbone NH of I78 (**Figure 2G**). The c36m simulations do not detect the same Cl⁻ binding geometries as the ones yielded by the AMOEBA simulations and experimental density. Furthermore, no alternative binding conformations within the neck region are detected for c36m simulations.

### The role of water in Cl⁻ permeation

Water plays a crucial role in ion channel permeation. The dimensions of the open neck state of bestrophin (**Figure 3A & 3B**) suggest the neck is sufficiently large to accommodate a fully hydrated Cl⁻ (radius ∼ 4 Å ^38^), the pore is wetted and permeable to ions. However, we observe that permeation through the neck occurs concurrently with the stripping of 1-2 water molecules from the first hydration shell of the ion (**Figure 3C**).

**Figure 3:**
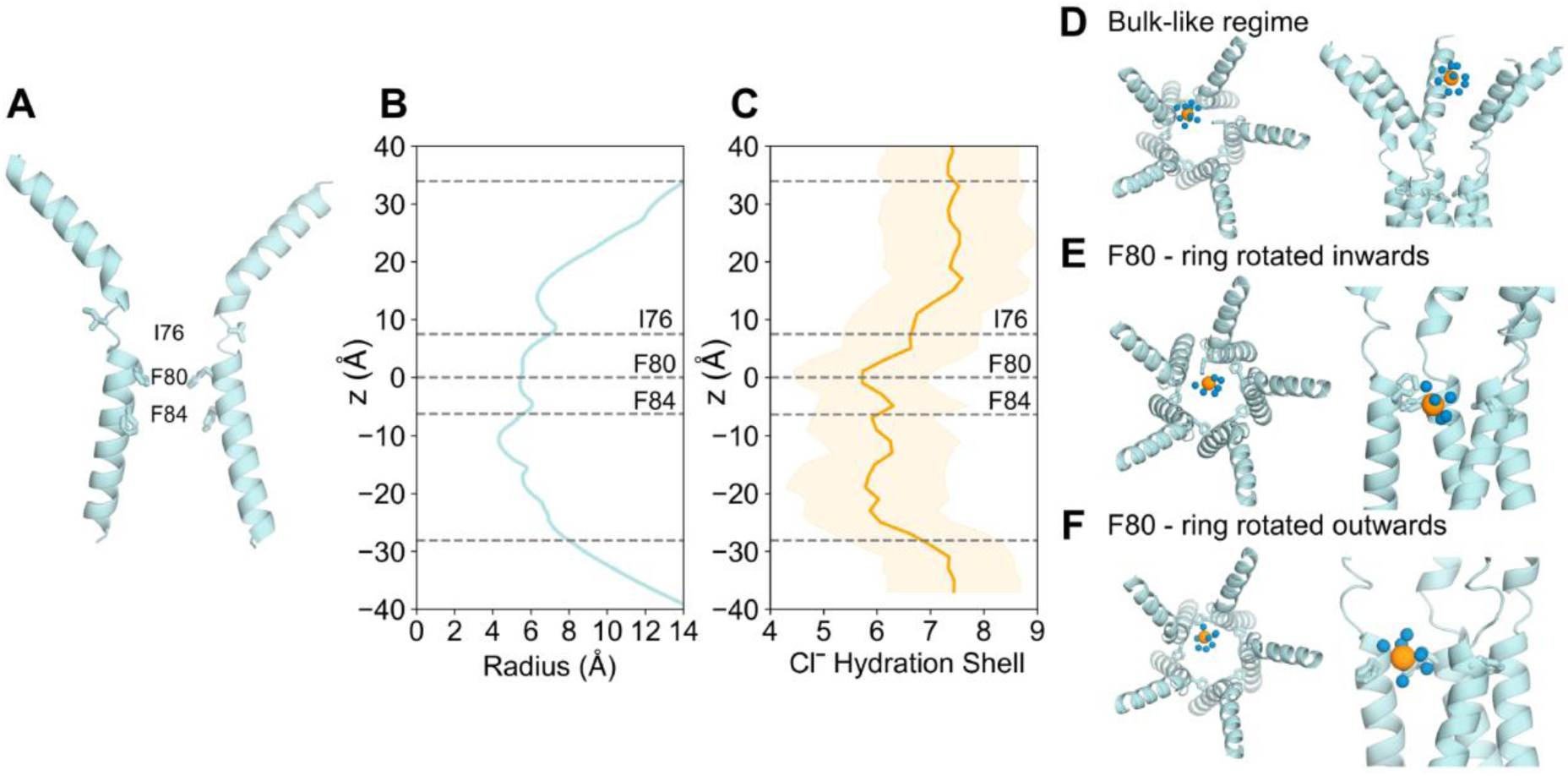
The role of water in Clˉ permeation in the open state AMOEBA simulations. **A** Cross-section schematic of the protein fragment of the open state bestrophin (PDB ID 8D1O) with the neck residues illustrated in licorice representation. **B** The pore radius profile of the open state fragment. The outer grey dashed lines represent the extent of the protein and the central lines indicate the *Z*-location of the neck residues where *Z* is the distance down the axis of the pore and *Z* = 0 corresponds to the *C*_α_ of F80. **C** The first hydration shell of Clˉ as a function of *Z*. The shaded orange region represents the standard error in hydration number at a given *Z*. Snapshots exemplifying the first hydration shell (blue spheres) of Cl⁻ (orange spheres) in **D** the bulk-like water regime within the protein fragment. **E** The partial loss of hydration shell due to anion-π interactions with the aromatic rings of F80 rotated inwards towards the pore axis. **F** The partial loss of Clˉ hydration due to anion-π interactions with the aromatic rings of F80 oriented outwards as in the experimental structure conformation.

The removal of water molecules allows for direct contact between Cl⁻ and the hydrophobic aromatic rings, mediated through anion-π interactions. The benzene ring of the phenylalanine has a large negative quadrupole moment that creates a partial negative charge on both faces of the π-system and subsequently a partial positive charge around the edge of the ring ^12,56,57^. As a result, the partial loss of hydration of Cl⁻ can be compensated by favorable interactions with these aromatic rings which could be regarded as “hydrophobic solvation” (**Figure 3E**). The aromatic sidechains of F80 possess rotational freedom allowing the ring to exist in two main conformations: flipped vertically inwards towards the pore axis (**Figure 3E)** or rotated outwards and in a flat orientation (**Figure 3F**). Both conformations facilitate hydrophobic solvation however note that the inwards flipped ring conformation occurs from a single protomer. There is no clear correlation between the flipping motion of sidechains and interactions with Cl⁻ in the immediate vicinity.

It has been hypothesized ^35–37,55^ that such a mechanism may mediate the stabilization of a dehydrated Cl⁻ at F80 and F84 through anion-π interactions based on the initial X-ray structure of chicken Best1 (cBest1), although this structure was later determined to be in the closed conformation. Mutagenesis studies of the cBest1 neck residues to alanine later showed no effect on ion selectivity and suggested that the role of anion-π interactions may reduce the energy barriers for Cl⁻ and other anions to permeate the neck region but not be the determining factor for charge selectivity ^58,59^. Other forms of hydrophobic solvation mechanisms have previously been observed in simulations of a biomimetic nanopore ^60^ and a hydrophobic protein binding site ^53^ whereby Cl⁻ moved through pores by partially dehydrating and forming energetically favorable interactions with hydrophobic contacts.

Importantly, these anion-hydrophobic interactions were only observed when electronic polarization was included in the molecular model and are not observed in equivalent simulations of the open state hBest1 when using fixed-charge (c36m) descriptions. We attribute this to the fact that anion-π interactions arise mainly from two factors: electrostatic interactions of the quadrupole moment and ion-induced polarization ^57^; the latter of which is not captured in c36m ^61^.

### Analysis of the partially open neck state

The partially open structure of hBest1 presents a more constricted neck region (**Figure 1E & 1G**). Critical conformational changes occur in residues F282, F283, and F276 concomitant with changes in neck residues I76, F80 and F84. I76 remains facing away from the pore axis as in the fully open state, while residues F80 and F84 and the hydrophobic gating apparatus adopt a closed-like conformation (**Figure 1G**) ^36^. In the experimental structure, an ion-like density is consistently captured at the edge of F84, even when processed with C1 refinement (Figure S1 D from ^36^) i.e., with no symmetry applied. This density could potentially represent a chemical constituent of the buffer system, in which Cl⁻ is the predominant ion, and so may represent a dehydrated Cl⁻. Furthermore, asymmetry in the neck can be observed as continuous movement between the closed and intermediate states in this region.

Trajectories of ions and water in the partially open state protein fragment (**Figure 4A**) see traces of Cl⁻ occupying locations just above F84 in the z-direction. This suggests that F84 is capable of accommodating Cl⁻; however, the pore is functionally closed as ions are not conducted across the neck, primarily due to the tight constriction formed by F80. Additionally, a small, transiently dewetted region can be found within the neck between residues S79 and F80, which reiterates that the partially open state is functionally closed (**Figure 4A**).

**Figure 4:**
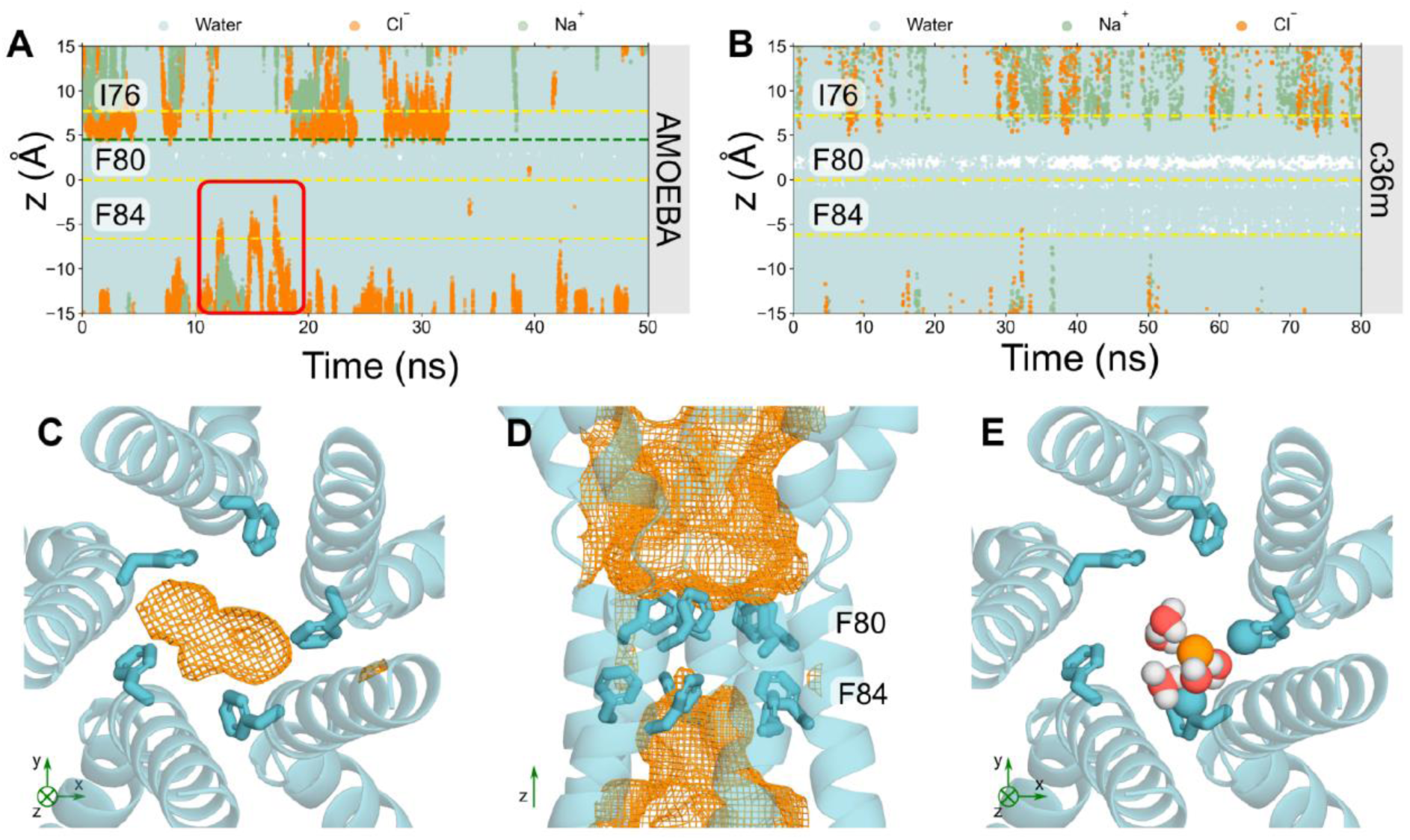
Analysis of the partially open state. Trajectories of water (cyan), sodium (green) and Cl⁻ (orange) ions in z-coordinates as a function of time within the neck region of the partially open state pore (PDB ID 8D1K) using **A** the AMOEBA forcefield or **B** c36m forcefield. The red box in **A** highlights a Cl⁻ that can occupy locations above F84 in the neck whereas nearly no ions can be seen in **B** with c36m. **C** Bottom-up view of F84 of the partially open state neck. The volume that Cl⁻ may occupy over the simulation is represented by the orange mesh. **D** Side-view of the neck region. F84 of the neck may accommodate for Cl⁻; however, the neck is not permeable to Cl⁻. **E** Bottom-up view of F84. Cl⁻ can be seen to interact with the aromatic ring through edgewise anion-π interactions. The coordinating atoms within the first solvation shell of the Cl⁻ (orange sphere) are represented as spheres.

The space that Clˉ can occupy is more clearly depicted in **Figures 4C & 4D**, where the orange mesh illustrates the volume of space that Cl⁻ occupies throughout the simulation. The majority of the neck region is devoid of Cl⁻ which could be associated with the dewetted region, since pore hydration acts as a precursor to ion permeation (**Figure 4D**). Mutagenesis studies have demonstrated that the conserved “IFF” motif of the neck acts as an effective gate, rather than a selectivity filter, because ionic selectivity remained constant in the mutant channel ^58,62^ which is in agreement with our results. F84 accommodates for a Cl⁻ through a similar mechanism to that seen in the previous section: the Cl⁻ partially dehydrates (**Figure S4**) to directly interact with up to 2 adjacent pore-facing phenylalanines simultaneously, mediated by their edge-wise interaction with the partially dehydrated anion (**Figure 4E**). Together, these observations support the provisional functional annotation of the channel as “partially open” i.e., it is indeed on-the-way to either opening or closing with the neck containing a consistently dewetted region and thus still presenting a barrier to ion permeation even though the neck (F84) can accommodate Cl⁻.

This barrier to ion permeation is not necessarily due to steric occlusion, but rather to the inability of the Cl⁻ to shed more of the first hydration shell. The partially open neck is wide enough to accommodate a dehydrated Cl⁻ (**Figure 4A & 4B**) but it is energetically unfavorable to lose more than ∼ 2-3 water molecules in the neck region at the tightest constriction which would be required for permeation and therefore ions do not pass through this region. For comparison, in the absence of polarization, c36m simulations suggest the neck is functionally closed and there is a significantly more prominent dewetted region between S79 and F80. F84 does not accommodate Cl⁻ but water can occupy this region (**Figure 4B**).

### Conclusions

We have performed MD simulations of the neck region of hBest1 channels in the open and partially open neck states using a fully polarizable forcefield (AMOEBA). Our results demonstrate that electronic polarization plays a key role in Cl⁻ accumulation at the helix dipole (the backbone nitrogen of F80) in the open state with significant overlap with an ion-like density previously assigned to a water molecule in the cryo-EM structure. Together, our data suggest that this location functions as a Cl⁻ binding site that contributes to anion selectivity in the open neck conformation of bestrophins. In contrast, fixed charge (c36m) simulations predict a different location for Cl⁻ accumulation that is not supported by experimental data, thus underscoring the importance of forcefield choice in simulation-based functional annotation of polarizable systems. Cl⁻ permeation through the open neck region is facilitated by edgewise anion-π interactions with the aromatic sidechains of F80 where, again, the influence of electronic polarization appears to exert decisive influence over the functional state of the channel. Simulations of the partially open neck state also support this permeation mechanism as neck dilation at F84 is able to accommodate Cl⁻; however, the pore remains functionally closed and is therefore “partially open”. Conversely, c36m simulations predict this 8D1K structure to be functionally closed. This study therefore provides new insights into bestrophin channel function although further studies are required to separate the individual contributions of structural elements to Cl⁻ permeation and selectivity within the ion pathway.

Overall, our results demonstrate the importance of modelling polarization in situations where polarizable moieties play an essential role in protein function. Appropriate treatment of electrostatics reveals more physically accurate behavior and provides new mechanistic insights into Cl⁻ permeation. MD simulations employing explicit polarization can therefore complement cryo-EM and other structural techniques in the identification and assignment of ambiguous non-protein densities such as anions.

## Acknowledgements

We thank T. Bertie Ansell for their insightful discussions regarding analysis and usage of PylipID. We would like to acknowledge the ARCHER2 UK National Supercomputing Service (http://www.archer2.ac.uk) and the University of Oxford Advanced Research Computing (ARC) facility in carrying out this work (http://dx.doi.org/10.5281/zenodo.22558). This work was supported by grants from the Biotechnology and Biological Sciences Research Council, the Engineering and Physical Sciences Research Council and National Institutes of Health grant R35GM149252.

## Author contributions

All authors designed research and wrote the paper. L.X.P performed research and analyzed data. S.J.T, J.C, M.S.P.S and T.Y obtained funding for the project.

## Supplementary Information

### Supplementary Methodological Details

#### Full protein embedded simulations

The full protein systems were prepared using a multiscale procedure. The protein is first coarse-grained (CG) and then embedded into a POPC bilayer and solvating with water and ∼ 0.5 M NaCl using Martini version 2.2 (1) and GROMACS 2021 package (2). This system is subject to 100 ns of equilibration in CG before being converted to atomistic representation with the CG2AT protocol (3). An equilibration period of 20 ns followed by a production run of 100 ns using the c36m forcefield (4) and mTIP3P water model was performed. The temperature was maintained at 310 K with coupling constant 1.0 ps by the Nosé-Hoover thermostat (5). Pressure was maintained at 1 bar with coupling constant 5.0 ps by the Parrinello-Rahman barostat (6). Short-range electrostatics were treated with the Verlet cutoff scheme at 1.2 nm cutoff and long-range electrostatics were treated with PME(7). C-alpha atoms were placed under harmonic restraints with a force constant of 1000 kJ/mol/nm^2^ to prevent the structure from deviating too much from the experimental structure.

**Figure S1:**
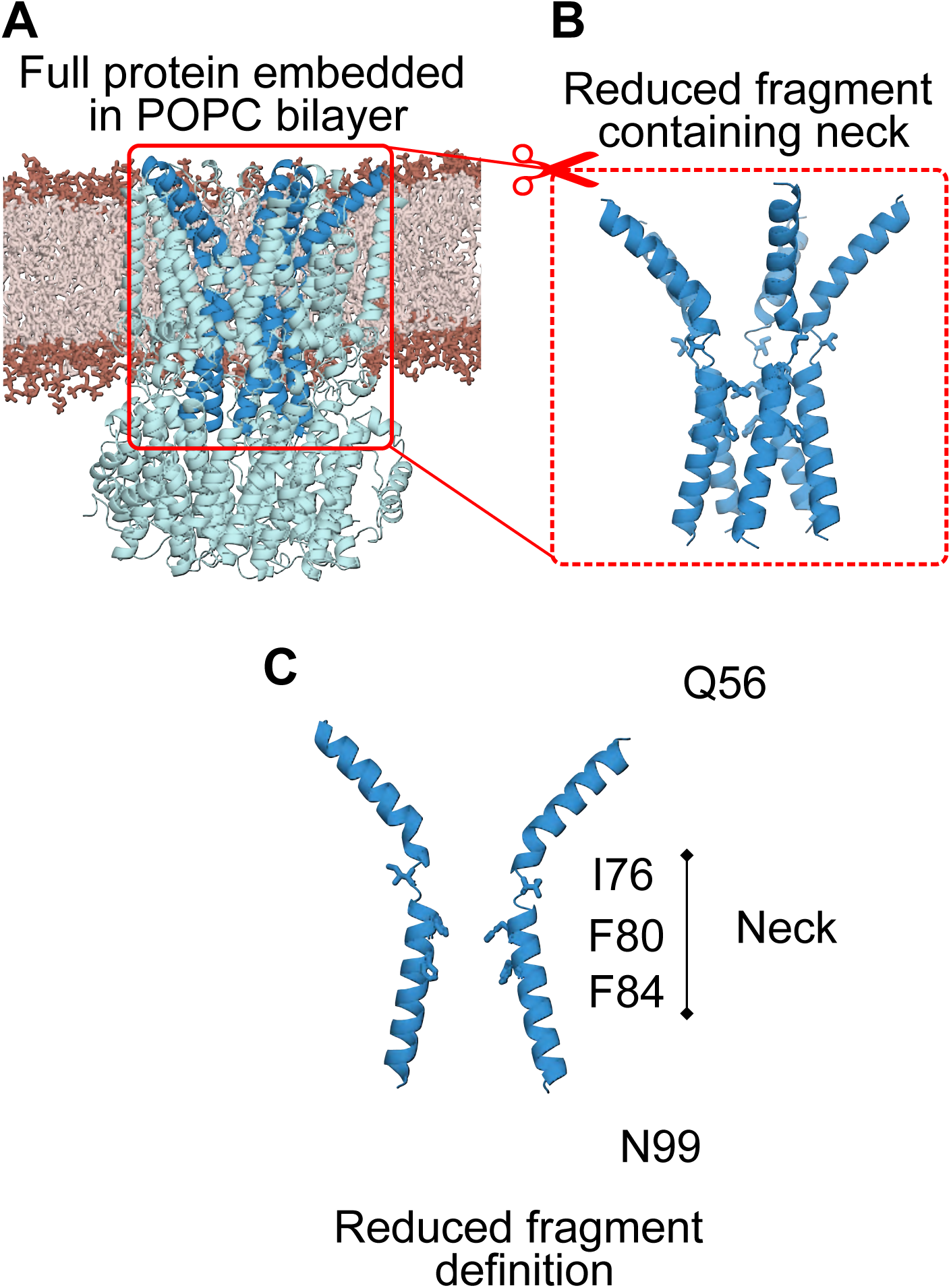
**A** Full protein (cyan) embedded in POPC bilayer (brown). **B** Protein fragment system used in simulations due to the high computational demands of the AMOEBA forcefield. The full protein embedded system in solution contains ∼200500 atoms. **C** The protein was truncated at Q56 and N99 such that the remaining fragment system contains the conserved hydrophobic neck region of interest (I76, F80, F84). The reduced protein fragment in solution contains ∼57000 atoms. Water and ions are omitted for clarity.

**Figure S2:**
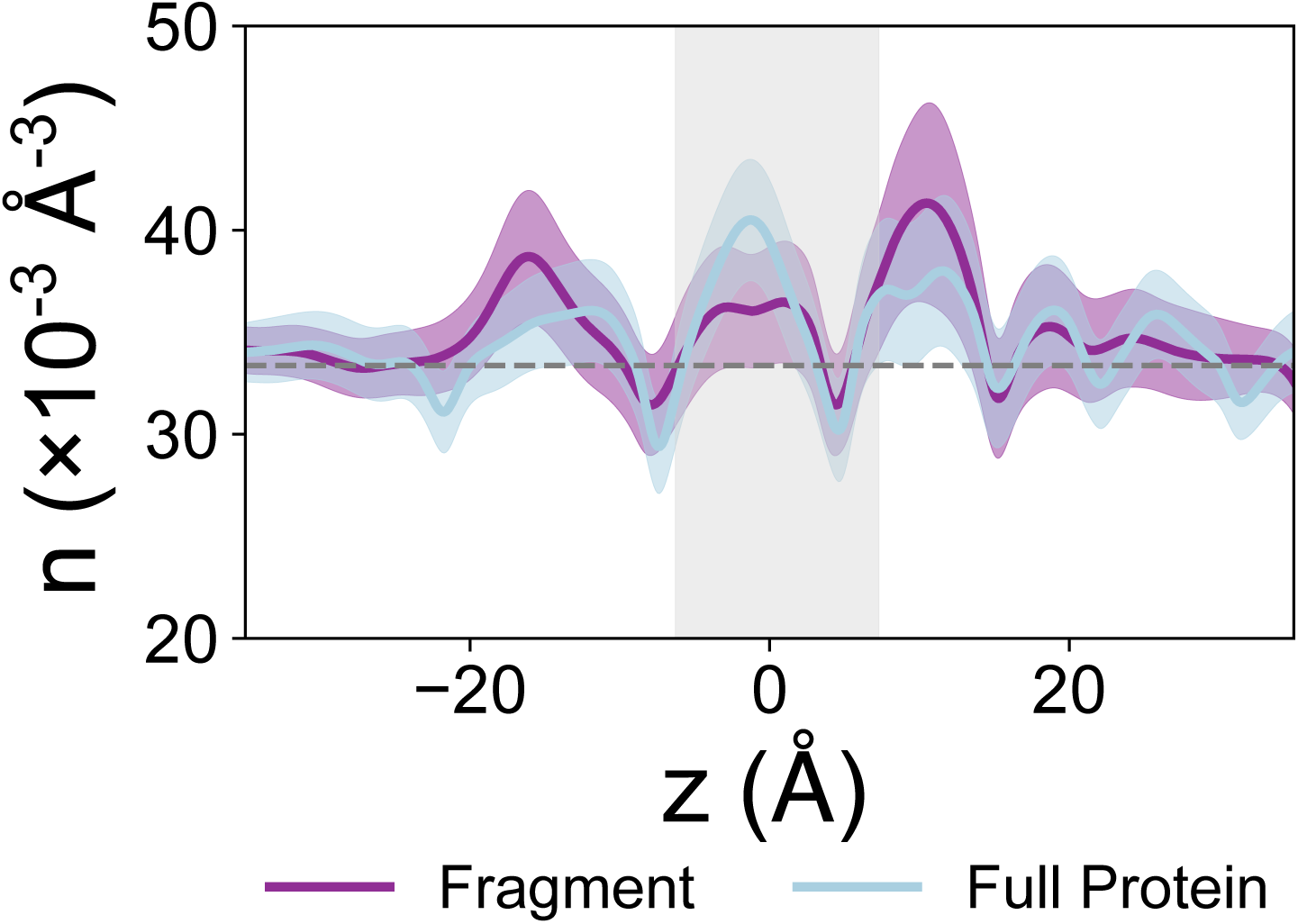
Protein fragment system validation. Time-averaged water density profiles within the pore for the full protein embedded system (light blue) compared with the protein fragment in solution (purple) using c36m. The shaded region represents the neck region, and the dashed grey line corresponds to the density of bulk water (33.37 nm^-3^). Confidence bands represent the standard deviation over the simulation.

**Figure S3:**
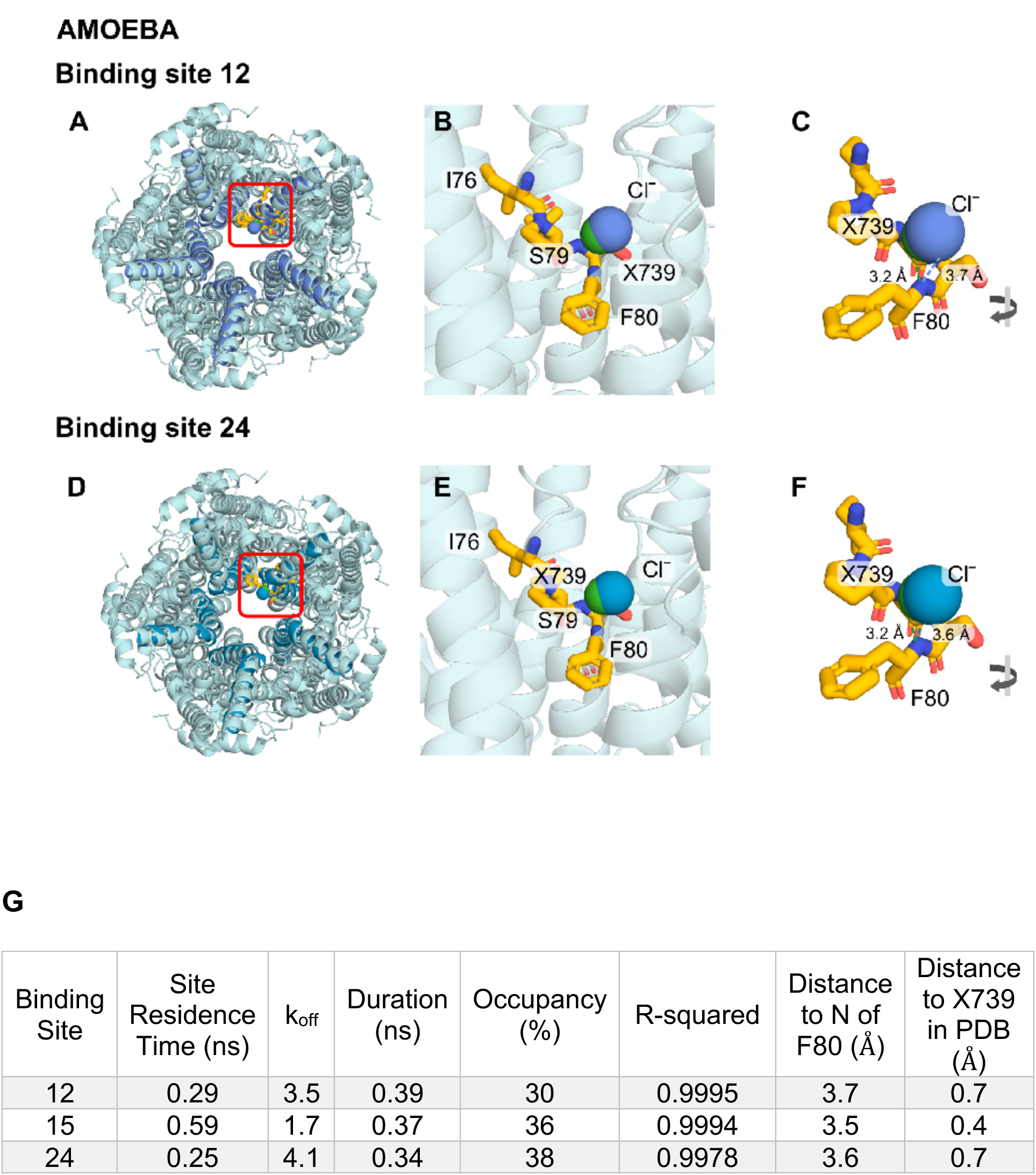
Alternative AMOEBA binding sites of the open state structure (PDB ID 8D1O). **A-C** Binding site 12 and **D-F** show binding site 24. Table **G** gives the binding site statistics where binding site 15 is the site shown in Figure 2 in the main manuscript.

**Figure S4:**
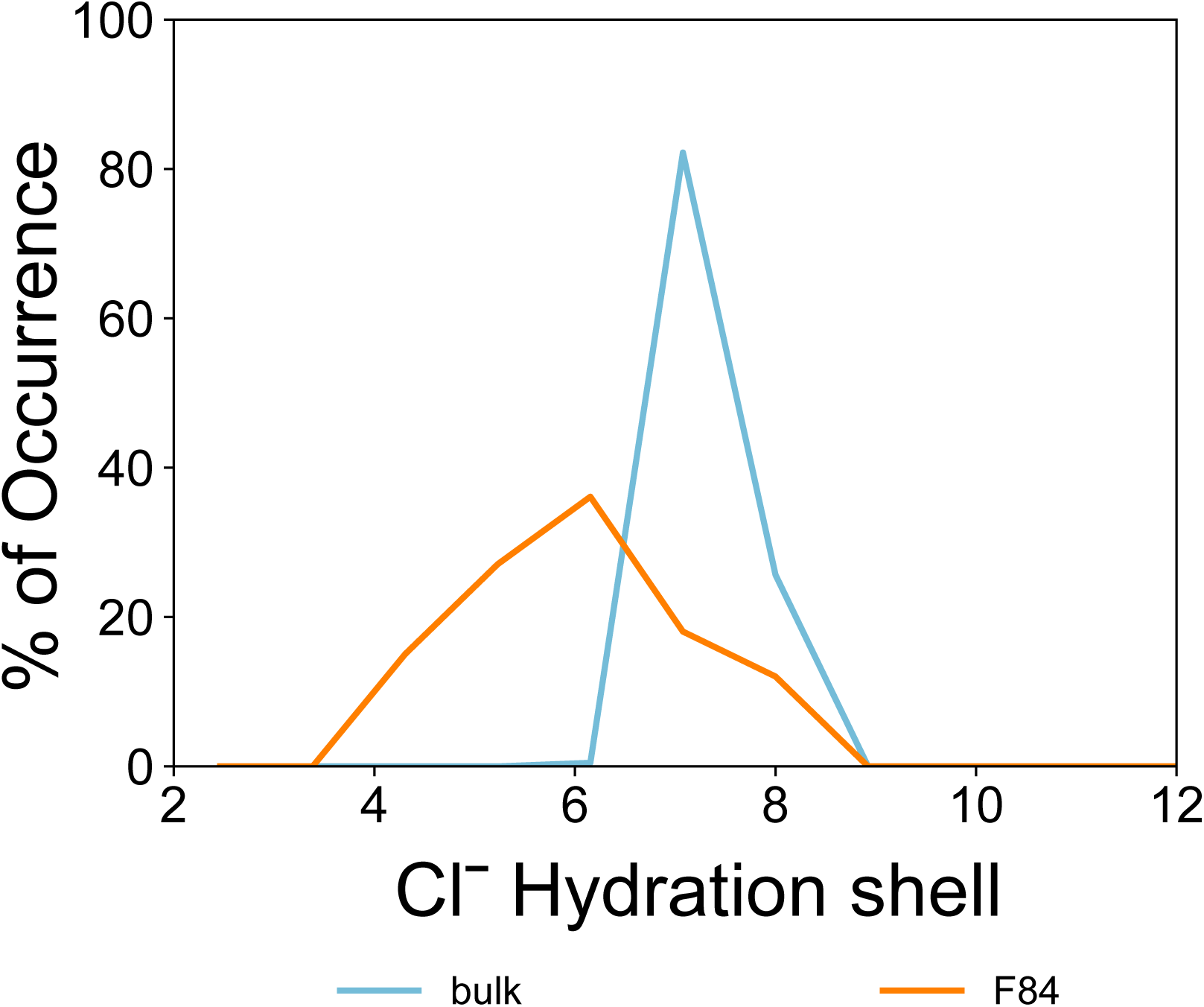
First hydration shell number of chloride ions in the partially open state structure (PDB ID 8D1K) in bulk (blue) compared with at the z-position corresponding to F84 (orange) within the pore in the AMOEBA simulation. Chloride loses 1-2 water molecules in its first hydration shell at F84 relative to bulk. This is comparable to the dehydration that occurs at F80 in the fully open state (PDB ID 8D1O).

